# Apelin signaling controls neuroendocrine homeostasis via Atoh8-dependent transcriptional regulation

**DOI:** 10.64898/2026.11.02.763639

**Authors:** Romain Menard, Anindita Neog, Caroline Halluin, Frederic Bonnet, Romain Madelaine

## Abstract

Apelin is a pleiotropic peptide hormone with established roles in cardiovascular function, angiogenesis, and metabolism, but its contribution to neuroendocrine development and function remains poorly understood. Here, using genetic loss-of-function and inducible gain-of-function approaches in zebrafish, we show that Apelin signaling is dosage-sensitive and required for normal development, reproductive capacity, and physiological homeostasis. Both *apelin* deficiency and *apelin* overexpression disrupted embryonic development and reduced adult lifespan, while exhibiting markedly reduced fertility in both sexes. Transcriptomic profiling revealed broad changes in neuroendocrine, hormonal, stress-response, and inflammatory genes with young adult mutant expression profiles resembling those of aged wild-type fish. Both loss and excess of Apelin also disrupted hypothalamic–pituitary–thyroid (HPT) axis development, reducing hypothalamic thyrotropin-releasing hormone-expressing cells, pituitary gonadotropic cell populations, and thyroid gland volume. Transcriptomic analyses identified *atoh8*, encoding a basic helix–loop–helix transcription factor, as a candidate effector of Apelin signaling. Apelin expression increased *atoh8* expression during early embryogenesis, whereas atoh8 mRNA partially rescued developmental defects in apelin mutants, including hypothalamic and pituitary markers, and thyroid volume. Conversely, *atoh8* crispants displayed thyroid and pituitary abnormalities, supporting a role for Atoh8 in neuroendocrine development. Together, these findings establish Apelin signaling as a dosage-sensitive regulator of zebrafish development and tissue homeostasis and identify Atoh8 as a functional downstream effector that partially mediates Apelin-dependent HPT axis formation. This work links peptide hormone signaling to neuroendocrine development and provides a framework for investigating its broader contributions to vertebrate physiology, reproduction and aging.

## INTRODUCTION

Apelin is a peptide hormone first isolated from bovine stomach extracts as the endogenous ligand of APJ (APLNR), a G protein–coupled receptor that had previously been an orphan receptor ^1^. Apelin is made as a prepropeptide precursor of 77-amino acid that is processed into several active isoforms, including apelin-36, apelin-17, and apelin-13 under the activity of the Furin enzyme ^2,3^. The system is conserved from fish to mammals, which points to a fundamental role in vertebrate physiology ^4^. Apelin and APJ are expressed in many tissues, including the heart ^5^, vascular system ^6^, skeletal muscle ^7^, brain ^8^, kidney ^9^, liver ^10^, lung ^11^, and adipose tissue ^12^. Across these tissues, apelin signaling coordinates development, energy metabolism, and tissue repair ^13–16^, through regulation of angiogenesis ^17,18^, myogenesis ^7,19^ and neurogenesis ^20^. Interestingly, Apelin/APJ expression has been associated with healthy aging ^7,13,18^, with anti-inflammatory ^21,22^, anti-fibrotic ^23,24^ and pro-regenerative properties ^13,16,19^. Finally, the Apelin pathway is also prominently active in endocrine organs, including the hypothalamus, pituitary, and the gonads ^1,25^ where recent studies point out a role in regulating reproduction ^26–28^. Altogether, the information above positions Apelin among the circulating hormones and systemic signals that connect peripheral and central tissue expression with neuroendocrine function regulating a broad variety of biological processes including development, reproduction, regeneration and aging.

Apelin and APJ expression decline with age across multiple organs ^29^, including the vascular ^18^, skeletal muscle ^7^, and the nervous system ^30,31^. This decline appears to be causal rather than incidental. Recent studies about skeletal muscle and vascular tissues beautifully illustrate this phenomenon ^7,18^. In skeletal muscle, contraction induces apelin production, and this production declines with age in mammals ^7^, while young mice lacking Apelin or APJ develop marked age-or disease-associated changes in tissue function ^19,32^. Skeletal fibers and plasma Apelin expression is reduced in aging and restoring apelinergic signaling promote regenerative myogenesis and counter age-associated muscle wasting ^7,32^. A parallel pattern is seen in the cardiovascular system. Mice lacking apelin or APJ show accelerated cardiovascular aging, whereas restoring Apelin in 15-month-old animals reduces cardiac hypertrophy and normalizes the exercise-induced hypertensive response ^29^. Plasma Apelin is likewise lower in elderly patients with peripheral artery disease, and in aged mice Apelin treatment restores blood flow recovery after ischemia by limiting endothelial senescence through APJ ^33^. Because apelin acts on both compartments, and because skeletal muscle repair depends on the endothelial niche ^19^, the apelinergic axis is well positioned as a shared upstream regulator of age-related decline in muscle and vessel. This convergence is most evident in sarcopenia, for which circulating apelin has been proposed both as an early diagnostic marker and as a therapeutic target ^7^.

In the central nervous system (CNS), Apelin–APJ signaling participates to maintain myelination of nerves, and both reduced oligodendrocyte APJ expression and decrease in circulating Apelin level with age impaired nervous system myelination ^31^. Pharmacological activation of APJ restores myelination *in vivo* in aged mice and increases *myelin* gene expression in human oligodendrocytes *in vitro*. This neuroprotective role extends to age-related neurodegenerative disease. Apelin peptides limit dopaminergic neuron loss and α-synuclein accumulation in MPTP models of Parkinson’s disease ^34,35^, and restrict amyloid-β production, tau phosphorylation and neuroinflammation in Alzheimer’s disease models ^36,37^. In ALS, apelin deficiency accelerates motor neuron loss and disease onset in SOD1^G93A^ mice ^38^. These studies indicate that failure of the Apelin–APJ axis contributes to the functional decline of the aging CNS ^31,38^. Supporting the above observations, both genetic deficiency and pharmacological blockade of Apelin signaling induce body-wide cellular senescence, at least partially, by activating senescence-promoting transcription factors ^13,29,39–42^. They also accelerate cardiovascular ^41^, skeletal muscle ^32^, renal ^9^, and reproductive ^43,44^ deficits in young mice. Altogether, these data indicate that Apelin sits at the intersection of systemic hormonal signaling and aging biology. Its progressive loss at once disrupts stem cell maintenance, promotes cellular senescence, and amplifies organismal aging ^7,29,31,45^, leading to degenerative phenotypes, including cardiovascular atrophy, micro-vessel rarefaction, and degenerative diseases such as sarcopenia and Alzheimer.

Apelin has also recently emerged as a regulator of reproduction in both sexes. Apelin and APJ are expressed throughout the hypothalamic–pituitary–gonadal (HPG) axis, including ovary, and testis of many species ^1,8,25–27,46,47^, where it acts in both an autocrine and paracrine manner ^1^. Most studies report that Apelin inhibits gonadotropin secretion: apelin-13 suppresses GnRH-stimulated LH release from the anterior pituitary in mice ^47^, and in pigs Apelin bidirectionally regulates GnRH- and Insulin-stimulated LH and FSH output through Akt, MAPK/Erk1/2 and AMPK signaling, with the direction and magnitude of the effect varying across the estrous cycle ^46^. This contrasts with the predominantly stimulatory actions of Apelin within the gonads, where it directly regulates steroidogenesis, cell proliferation, apoptosis and sexual hormone secretion ^1^, suggesting that the apelinergic system acts in opposite directions at different tiers of the axis. Beyond the gonads, Apelin/APJ regulates placental proliferation, apoptosis, angiogenesis in mammals ^48,49^ and disruption of these processes is associated with pregnancy complications in humans ^49^. Disturbed Apelin signaling has been linked to reproductive disorders such as polycystic ovary syndrome, endometriosis, and ovarian cancer, which has drawn interest in the apelinergic system as a therapeutic target for infertility ^1,50^. Experimental manipulation of Apelin/APJ signaling directly alters reproductive function. In male rats, central or peripheral Apelin-13 administration suppresses LH and testosterone and reduces Leydig cell number ^51^, whereas pharmacological inhibition of APJ in juvenile mice impairs testicular proliferation, increases apoptosis, and reduces androgen receptor expression ^27,52^. In diabetic mice and cultured human testes, aberrant APLN/APJ activation disrupts the blood–testis barrier, and its inhibition restores spermatogenesis ^53^. In women, defective Apelin/APJ signaling in granulosa cells derived from Turner syndrome patients impairs cell-cycle progression, which is rescued by Apelin supplementation ^54^, and in infertile men, higher sperm Apelin levels correlate with abnormal sperm morphology ^55^. Together, these findings indicate that both excess and insufficient Apelin signaling compromise reproduction, suggesting that tightly regulated Apelin levels are required for reproductive competence. Apelin is therefore one of the few circulating factors with documented roles in both systemic aging ^56^ and reproductive longevity ^45^.

As mentioned above, the Apelin/APJ system is expressed throughout the HPG axis, and decline with age ^29,57^. Interestingly, Apelin signaling has also been reported to influence thyroid gland function via modulation of the hypothalamic–pituitary–thyroid (HPT) axis ^25,58^, and the apelinergic pathway may be an underappreciated regulator of neuroendocrine signaling during development and aging. Supporting this hypothesis, it has been recently reported in patients with acute, severe hypothyroidism after thyroidectomy, that Apelin expression level in the serum dropped markedly ^59^. Despite these associations, it remains unknown whether the decline in systemic Apelin expression contributes to HPT axis developmental defects or age-related remodeling.

The HPT axis is a neuroendocrine system highly sensitive to age-related hormonal change ^57,60^ and variations in thyroid hormones (TH) expression appear to occur similarly in men and women during aging ^61^. In women, thyroid hormones regulate menstrual function, fertility, and the maintenance of pregnancy ^62,63^, while thyroid autoimmunity is associated with infertility and low ovarian reserve in subsets of women ^63^. Interestingly, the thyroid directly connects to ovarian aging with TH signaling, including local T3 metabolism within the ovary, being implicated in preserving the functional ovarian reserve during aging ^63,64^. In men, hormonal thyroid dysregulation alters sex steroids and hormone-binding globulin, affecting sperm function and reproduction ^62^. The testicular development is also highly sensitive to thyroid hormone activity with the proliferation and differentiation of Sertoli, Leydig, and germ cells regulated by the HPT axis ^65–67^. The above evidence indicates that the HPT and HPG axes are mutually linked throughout life ^61^, and that age-related hormonal change, such as Apelin expression, are well positioned to influence both reproduction and aging.

In the neuroendocrine HPT axis, hypothalamic thyrotropin-releasing hormone (TRH) stimulates pituitary thyroid-stimulating hormone (TSH), which drives the thyroid to secrete thyroxine (T4) and triiodothyronine (T3) ^68,69^. Through these several layers of control, the HPT system integrates diverse aspects of organismal physiology, including metabolic rate, stress responses, fertility and longevity, and undergoes characteristic morphological and functional changes with age ^70^. During aging, thyroidal iodine uptake falls, synthesis of free T4 and free T3 declines, and reverse T3 (the inactive product of T4 deiodination) rises while serum TSH generally stays within the normal range ^70^. Because optimal thyroid function is considered a prerequisite for normal reproductive capacity ^62^ and healthy aging ^70^, we investigated the role of Apelin in the homeostatic maintenance of the HPT axis.

To investigate this relationship, we leveraged the zebrafish model, which shares a conserved neuroendocrine architecture with mammals ^71–74^. Using this model organism, we investigated how Apelin signaling affects development, aging and reproduction. Loss- and gain-of-function Apelin zebrafish models display overlapping developmental defects and aging signatures, including growth defects, fertility impairment, elevated stress responses, and shortened lifespan. Transcriptomic profiling during development and aging revealed aging signatures both dependent and independent of Apelin that are conserved across tissues. These observations converge on physiological processes known to be under neuroendocrine regulation. In seeking molecular effectors downstream of Apelin signaling, we established Atoh8 (also known as Math6) as a transcription factor being negatively and positively regulated by loss- and gain-of-function respectively. Atoh8 is a member of the atonal subfamily of basic helix-loop-helix (bHLH) transcription factors, a superfamily of regulatory proteins with critical roles in cell fate determination, differentiation, and development across diverse tissues ^75–77^. Unlike some bHLH factors that serve as master regulators of specific cell lineages, ATOH8 fulfills broader developmental roles^76,78,79^ with established functions in skeletogenesis, myogenesis, neurogenesis, and cardiovascular development ^78–81^, processes that overlap extensively with those regulated by the Apelin peptide. Supporting this observation, Atoh8 expression can significantly reverse phenotypic defects observed in the Apelin mutant, including development of the HPT axis. Together, these findings establish Apelin as a regulator of HPT homeostasis and identify Atoh8 as a previously unrecognized molecular effector of Apelin signaling, revealing an Apelin–Atoh8 axis with potential therapeutic relevance for metabolic-endocrine disorders characterized by systemic hormonal and thyroid dysfunction.

## RESULTS

### Loss of *apelin* function leads to growth and reproductive defects

Apelin is a peptide hormone expressed throughout the body whose levels decline with age. To better understand the role of Apelin in development and aging, and assess its physiological functions, we previously developed and reported an *apelin* knock-out model generated using the CRISPR/Cas9 system ^82^. We observed developmental phenotypes in ∼40% of embryos at 24 hpf (hours post-fertilization) issued from an incross between heterozygous *apelin* mutants (Fig. 1A, B), suggesting that both heterozygous and homozygous genotypes can affect embryonic development. In addition, we identified that the hatching of these larvae from the chorions was strongly reduced at 3 dpf (days post-fertilization) with ∼75% of unhatched larvae (Fig 1C). From 3 dpf onward, distinct phenotypes emerge (Fig. 1D, E), with some affected larvae harboring reduced body size (mild phenotype), including brain and retina, and others also developing cardiac edema (severe phenotype). Until 5 dpf, the survival of larvae from heterozygous mutant incross is similar to larvae obtained from wild-type siblings (Fig. S1A). To obtain an assessment of larval and juvenal survival of the *apelin* mutant until adulthood, we determined mortality and genotype of 5 dpf larvae until they reach sexual maturity (Fig. 1F). When larvae were screened based on the mild phenotype at 5 dpf, we identified that larvae survived until adulthood, and that the large majority of the adults genotyped by PCR have one or two copies of the mutation, suggesting that the mild phenotype can be associated with the *apelin* mutation, either heterozygous or homozygous. However, when larvae have not been screened on the phenotype, we quantified that ∼20% of the larvae issued from heterozygous incross will not survive until adulthood, indicating that larvae carrying a cardiac edema (severe phenotype) will most likely not reach the adult stage. Among the surviving others, we identified adult wild-type, heterozygous and homozygous *apelin* mutant fish by genotyping, confirming that both heterozygous and homozygous mutants can reach the adult stage. Interestingly, both female and male adult mutants show a reduction in body size with the adult homozygous mutants showing a stronger phenotype (Fig. 1G, H), suggesting that heterozygous mutant can partially compensate for the early growth defect observed between 1 and 5 dpf. Finally, we assessed the reproductive performance of the young adult (< 6 months) heterozygous or homozygous *apelin* mutant fish and revealed a dramatic reduction in fertility in the male and female homozygous mutant fish (Fig. S1B). In rare situations (three times), we obtained eggs from homozygous female mutants crossed with wild-type males, but only a small percentage of the eggs were fertilized (Fig. S1C). Given this constraint about using homozygous mutant for reproduction, heterozygous carriers for the *apelin* mutation were used for all subsequent analysis in this study. Altogether, the phenotypic analysis of the *apelin* mutants, indicates that this peptide hormone controls growth and reproduction in zebrafish.

**Figure 1.**
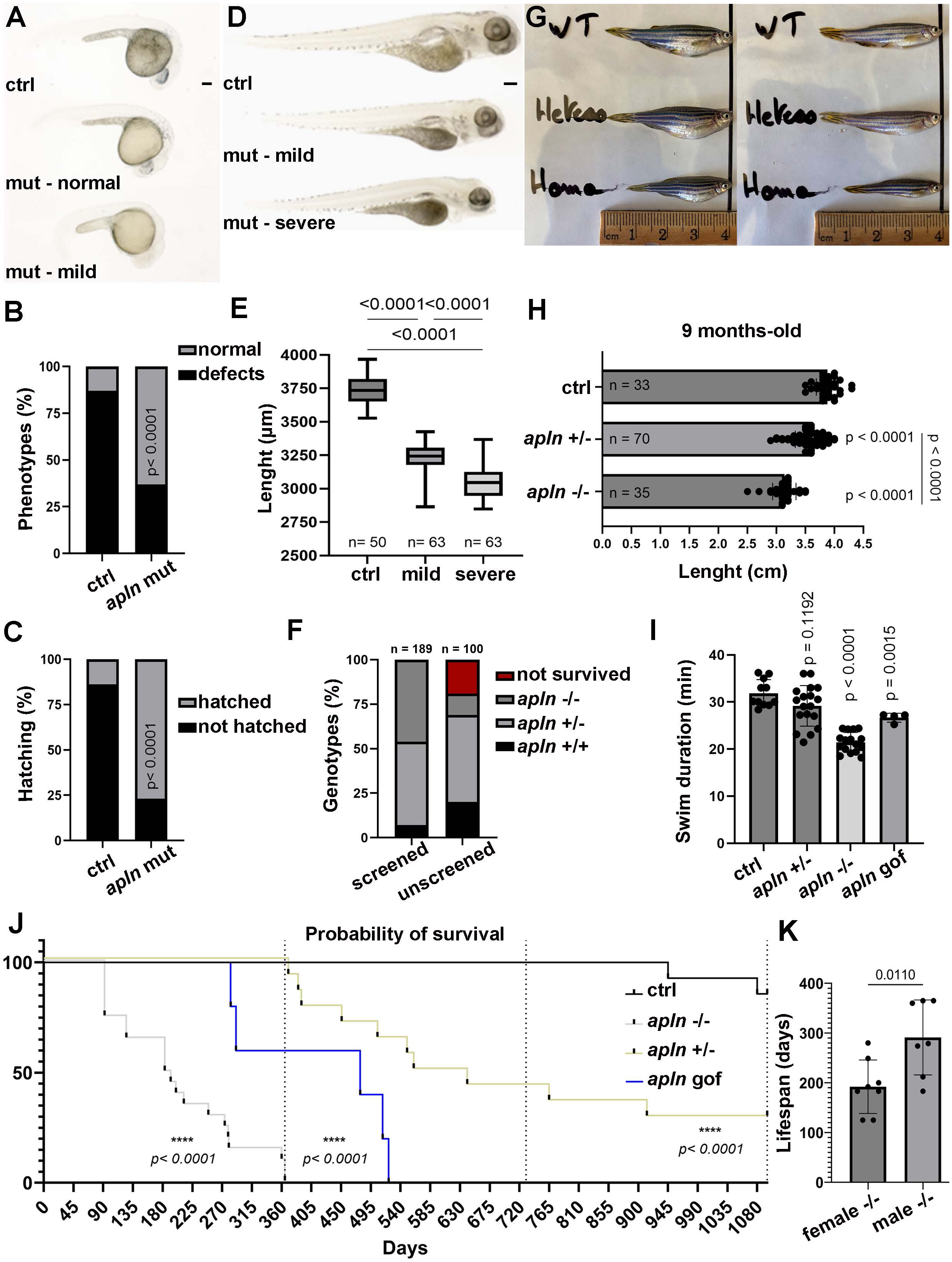
*apelin* zebrafish mutants have growth defects and shorter lifespans. **(A)** Representative images of *apelin* mutant larvae at 1 dpf showing either normal or mild phenotypes. **(B)** Quantification of control (n=100) vs *apelin* mutants (n=100) larvae at 1 dpf with normal development (black bars) or developmental delays (grey bars). Significance is determined by Fisher’s exact test, two-sided, contingency table. **(C)** Hatching analysis of control (n=175) vs *apelin* mutants (n=78) larvae that have hatched (black bars) or not hatched (grey bars) at 3 dpf. Significance is determined by Fisher’s exact test, two-sided, contingency table. **(D)** Representative images of *apelin* mutant larvae at 5 dpf showing body size reduction (mild) and oedema (severe) phenotypes. **(E)** Body size at 5 dpf of control (n=50), *apelin* mutants mild (n=63) and severe (n=63) phenotypes. Significance is determined by Mann-Whitney test, non-parametric. **(F)** Distribution representation of *apelin* mutant fish reach adult stage after being screened on phenotype (left graph; n=189) or unscreened (right; n=100) at 5 dpf. Genotyped identification wild type (black), heterozygous (light grey), homozygous (dark grey), dead fish (red). **(G)** Representative images of *apelin* mutant adult length at 9-month-old; wild-type (top), heterozygous (middle) and homozygous (bottom) fish; female (left side) and male (right side). **(H)** Body size quantification of wild-type (n=33), heterozygous (n=70), and homozygous (n= 35) *apelin* mutant fish. Significance is determined by Mann-Whitney test, non-parametric. **(I)** Analysis of the swimming performance of wild-type (n=10), *apelin* mutant heterozygous (n=18), *apelin* mutant homozygous (n=17), and apelin overexpression (n=4) at 9-month-old. Significance is determined by Mann-Whitney test, non-parametric. **(J)** Survival analysis over a 3-year period of wild-type (n=8), *apelin* mutant heterozygous (n=14), homozygous (n=20) siblings and *apelin* overexpression (n= 5). Significance is determined using Kaplan-Meyer simple survival analysis. **(K)** *apelin* mutant homozygous female (n=8) and male (n=7) gender lifespan analysis. Significance is determined by t-test, two-sided, unpaired. Error bars represent standard deviation (s.d.). Scale bars: 100 μm.

### The proper level of *apelin* expression is important for normal embryonic development

To better understand the role of Apelin, we adapted a previously established gain-of-function strategy ^83^, replacing Atrogin-1 with Apelin in the Tg(*ubi:Lox-stop-Lox*) cassette. A muscle specific driver was selected because the Apelin prepropeptide is largely and naturally expressed, processed and secreted into the circulation from this tissue to reach organs throughout the body in a systemic manner ^7,16,32^. The expression of *apelin* is temporally controlled by 10 µM of hydroxytamoxifen (4-OHT) or tamoxifen (TAM) treatment in larvae or adult respectively. To further characterize these phenotypes, we again defined three groups: normal (no morphological defects), mild (growth defect) and severe (also include pericardial oedema). We followed phenotype development between 3 and 5dpf from wild-type siblings, control larvae treated with 4-OHT, larvae from heterozygous *apelin* mutant incross and larvae overexpressing *apelin* (gain-of-function; gof between 3 and 5 dpf) (Fig. S1D). To determine whether phenotypic severity is associated with genotype, larvae were sorted by phenotype at 3 dpf, and each group was followed individually until 5 dpf, with each larva being genotyped at the terminal time point (see legend of Fig. S1D for details). In *apelin* mutant at 3 dpf, larvae were distributed across the three groups as follows: normal (20.88%), mild (53.82%), and severe (23.82%) phenotypes, with correlating genotype frequencies of wild-type (21%), heterozygous (51%), and homozygous (28%). Between 4 and 5 dpf, we observed shift away from the normal phenotype, with increasing numbers of larvae transitioning toward a mild or severe phenotype. Genotyping of single larvae revealed that the majority undergoing this phenotypic progression were heterozygous (65.26%) or homozygous mutants (14.67%). Across all time points, larvae of both heterozygous and homozygous genotypes were found in all three phenotypic groups, as well as a small number of wild-type individuals (Fig. S1D). Notably, heterozygous larvae were overrepresented in the mild group (52%) and almost equally distributed between the normal (44%) and severe groups (55%), while homozygous larvae were predominantly found in the mild (30%) and severe (40%) groups, with only a minority classified as normal (13%). This observation suggests that embryos that did not survive until 5 dpf were most predominantly homozygous mutants. In contrast, when wild-type clutches were subjected to the same classification scheme, approximately 6% of larvae showed early phenotypes but the majority recovered to a normal morphology at 5 dpf (Fig. S1D). Importantly, phenotypic progression was strictly unidirectional: *apelin* mutant fish could advance from normal to mild or from mild to severe, but no regression to a less severe phenotype was observed. Using the overexpression strategy with 4-OHT treatment at 10 µM at 3 dpf for 48 hours, we followed larvae overexpressing *apelin* at day 4 and 5 of development. Phenotypes onset was visible as early as 24 hours after treatment initiation with the appearance of cardiac edema being a predominant feature. Wild-type control larvae treated with 4-OHT showed neither edema or growth defects; but mild toxicity, with approximately 5% of larvae dying within 48 hours of treatment, a rate similar to that of the treated gof larvae (Fig. S1D). Together, the above analysis indicates that both loss and gain of function of Apelin signaling affect tissue growth and the formation of cardia edema, suggesting that the expression of the *apelin* gene has to be tightly controlled for the proper regulation of embryonic development.

### Dysregulation of Apelin activity affects lifespan

Given the presence of pericardial edema in the functional analysis and the established role of Apelin in regulating cardiac function ^84,85^, we next measured heart rate across all phenotypic subgroups. In both the loss- and gain-of-function lines, heart rate was markedly reduced relative to controls. Control larvae maintained a heart rate of approximately 180 bpm, while mild-phenotype larvae averaged 150 bpm and severe-phenotype larvae averaged 100 bpm, implicating cardiac dysfunction as a shared feature of both loss and gain of function models (Fig. S1E). We also assessed locomotion in zebrafish larvae between 5 and 7 dpf in *apelin* mutant (mild phenotype) and overexpression conditions and identified reduced locomotor capacity in both loss and gain of Apelin function (Fig. S1F). Next, we investigated the effects of the loss and gain of function strategy in adult fish. Similarly to the adult homozygous mutant phenotype, adult fish with *apelin* gain-of-function induced by tamoxifen treatment (TAM) once a week for 24h starting at 1 month-old, harbor a reproductive deficit, supporting the previous observation that both reduced and increased expression of *apelin* can have detrimental effects on the reproductive system (Fig. S1B). In addition, we assessed swimming capacity in adults, and observed that similarly to larvae, both homozygous *apelin* mutant and gof conditions showed reduced performance at 9 months-old (Fig. 1I). However, whether the reduced locomotor activity observed in larvae and adults is dependent on defects in muscle and/or cardiovascular function remains undetermined. Finally, we monitored the lifespan of adult fish over a 3-year period to determine whether Apelin dysregulation can affect longevity (Fig. 1J). Both the loss- and gain-of-function genotypes exhibited shortened lifespans relative to wild-type. Within the homozygous cohort, 50% mortality was reached by 180 dpf and no individual survived past 1 year. Among heterozygous fish, only 30% reached the 3-year endpoint. Notably, homozygous females had an average age at death of approximately 6 months compared to approximately 9 months in males, suggesting a sex-differential effect of Apelin signaling, with a more severe impact on females (Fig. 1K). Fish overexpressing *apelin* survived slightly longer compared to homozygous mutants, with 50% mortality at 495 days, though complete mortality was reached by 18 months. Taken together, these results demonstrate that both *apelin* loss and gain of function are deleterious for the homeostasis of the body when not properly controlled, leading to growth defects, early aging phenotypes and death.

### Apelin regulates the expression of genes associated with reproduction and aging signatures

To further investigate the molecular mechanisms underlying *apelin*-deficient dependent aging, we performed bulk RNA sequencing on dissected brain, eyes, and trunk tissue from adult *apelin* homozygous mutant and wild-type control siblings fish at 12 months of age, as well as wild-type aged fish at 48 months of age. Four fish per group (two male, two female) were used, yielding mRNA expression profiles that enabled three pairwise comparisons per tissue: wild-type versus mutant at 12 months, 48-month wild-type versus 12-month mutant, and 12-month wild-type versus 48-month wild-type — resulting in nine dysregulated gene lists in total (Fig. 2A and Table S1-9). The largest transcriptomic divergence was observed between wild-type and *apelin* mutant fish at 12 months (Fig. 2B). Notably, the differential level of gene expression between 12-month-old mutants and 48-month-old wild-type fish was comparatively smaller, suggesting a pattern of accelerated aging in the mutant (Table S10-12), an observation correlating with their reduced lifespan (Fig. 1J, K). To better characterize the aging signature dependent on Apelin signaling, we intersected the three comparison lists within each tissue (Fig. 2C), generating three Venn diagrams that allowed us to distinguish genes regulated by Apelin that are independent or associated with aging. This approach identified 403, 483, and 419 aging-associated and 536, 382, and 306 aging-independent genes regulated by Apelin in the trunk, eyes, and brain respectively (Fig. 2C). Next, to identify genes commonly dysregulated across all three tissues, we then intersected these per-tissue lists, yielding two final gene sets representing the shared aging signature across brain, eyes, and trunk: 114 genes and 83 genes dysregulated in an Apelin-dependent manner that are associated with or independent of aging signature respectively (Fig. 2D). Among the genes associated with aging, some were similarly dysregulated across all three tissues, such as *klf9*, *fkbp5*, *scg5*, *scamp4*, *sulf2a, vegfba* and *vegfaa*, supporting a role for Apelin signaling in angiogenesis, peptide secretion and stress response. Of these, *fkbp5* and *klf9*, both upregulated, are well-established glucocorticoid-responsive genes, and *klf9* is also a direct thyroid hormone target ^86,88,90^, suggesting that Apelin signaling dysregulation alters glucocorticoid and possibly thyroid hormone signaling. In particular, we identified by GO a set of genes, dysregulated in the *apelin* mutant and associated with aging, converging on neuroendocrine biology such as hormone transport and secretion (Fig. 2E, Tables S13-15). Among genes regulated in an aging-independent manner by Apelin, the GO enrichment identified gonad development and reproductive behaviors, as well as regulation of inflammation as some of the main dysregulated processes (Fig. 2E, Tables S16-18). Strikingly, the *mnd1* gene, which is essential for meiotic recombination and is associated with neurological and reproductive defects, is downregulated ∼141x in the *apelin* mutant. Its human ortholog is linked to primary ovarian insufficiency and azoospermia ^87,89^, suggesting a potential role for the Apelin peptide in gametogenesis. We also identified genes and GO terms dysregulated in the brain of *apelin* knockout fish that are aging-associated or independent (Fig. S2B and table S19-20) and could underly reproductive and aging deficit observe in the *apelin* mutant. This analysis reveals a subset of brain-specific expressed genes regulated by Apelin with direct relevance to HPT axis homeostasis. In particular, genes involved in the thyroid hormone signaling such as *nr1d1* and *nr1d2* (Rev-erbα and Rev-erbβ) are upregulated. In mammals, *NR1D1* is encoded on the antisense strand of the *THRA* locus ^91–93^, and *Rev-erbα/β* proteins are core circadian repressors whose activities regulate the rhythmic secretion of TRH and TSH ^94,95^. Interestingly, one of the core regulators of the circadian clock, BMAL1 that heterodimerizes with CLOCK to drive rhythmic transcription, is strongly downregulated in our dataset. In mammals, *Rev-erbα/β* directly represses *Bmal1* transcription ^96–98^, an observation consistent with *nr1d1* and *nr1d2* upregulation in the brain transcriptome, and a potential dysregulation of HPT signaling. *BMAL1* has an established role in HPT-dependent regulation of reproduction. Knockout mice for *BMAL1* have defective hormone secretion and signaling, and are infertile, with impaired reproductive organs and gametes development ^99,101^. Finally, biological processes network analysis reveals clusters of interactions indicating that hormonal and inflammatory responses are the main component dysregulated in aging-dependent and independent manner respectively (Fig. S2C). Altogether, the convergence of transcriptomic and phenotypic evidences supports a role for Apelin signaling in controlling reproductive ang aging processes, potentially via the regulation of HPT axis activity.

**Figure 2.**
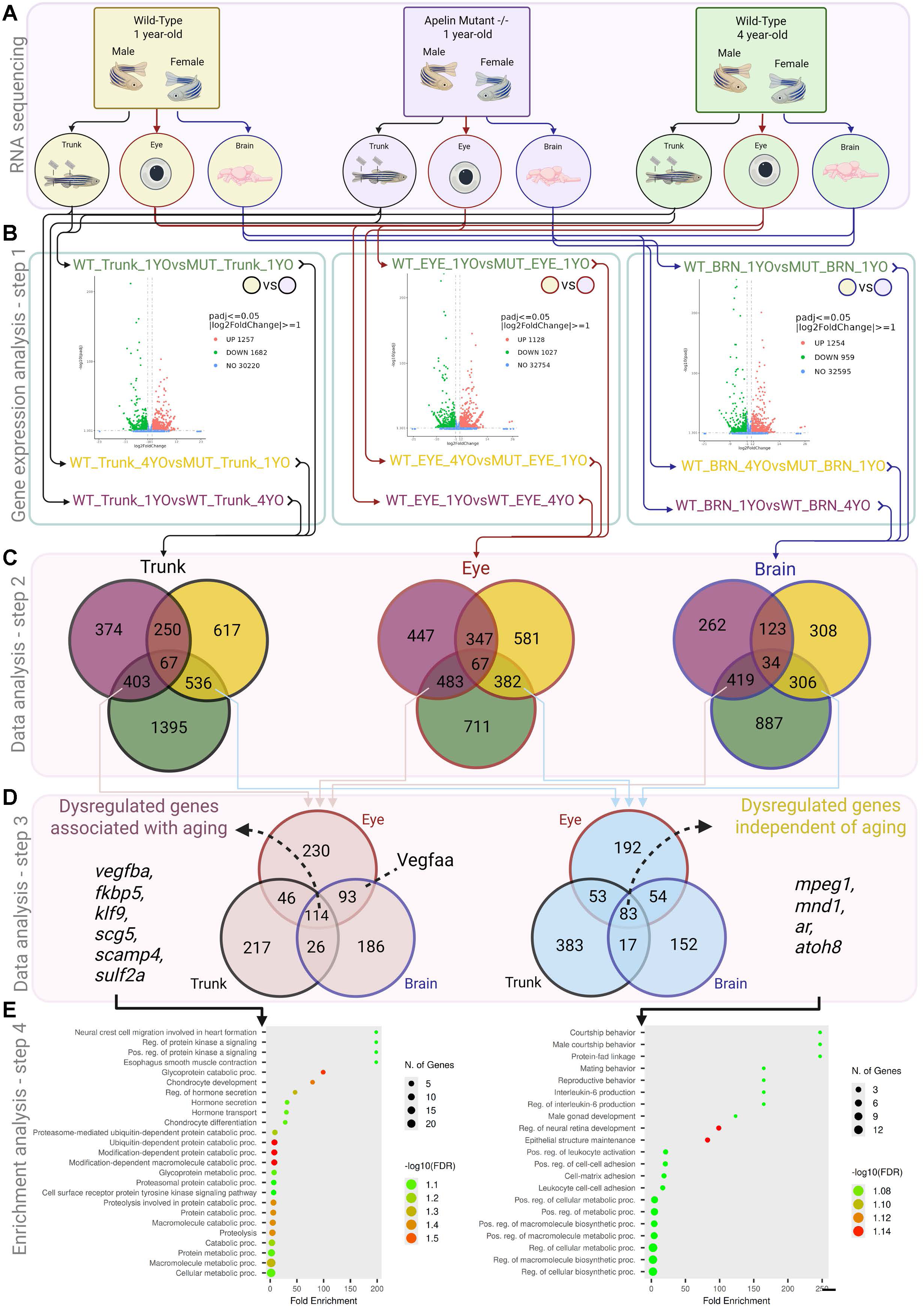
Identification of aging-dependent and aging-independent gene expression signatures associated with Apelin expression. **(A)** Each fish (n=4 per condition) was dissected, and the brain, eyes, and trunk were isolated. Each condition is represented by a background color: wild-type 1-year-old (yellow), *apelin* mutant 1-year-old (blue), and wild-type 4-year-old (light green). Each tissue is denoted by a distinct border color and corresponding arrows: trunk (black), eyes (red), and brain (dark blue). This scheme allows each tissue to be tracked across conditions and illustrates how the comparisons were made. **(B)** Volcano plots of downregulated (red) and upregulated (green) genes in the 3 different adult tissues between experimental conditions**. (C)** The gene expression analysis comprised three lists per condition and three lists per tissue, yielding nine sets of dysregulated genes. The gene lists for each tissue were then intersected (indicated by the black, red, and blue arrows), producing three sets of gene lists and corresponding Venn diagrams—one per tissue—identifying the dysregulated genes shared between WT_1YO vs. Apelin_Mut_1YO, WT_1YO vs. WT_4YO, and WT_4YO vs. Apelin_Mut_1YO. From these intersections, two gene sets were isolated: Apelin-dysregulated genes associated with aging (pink) and Apelin-dysregulated genes independent of aging (light blue). **(D)** Aging-associated (mauve) and aging-independent (green) gene sets from the trunk, brain, and eyes were intersected to reveal genes dysregulated in a similar manner across all three tissues. **(E)** GO (Gene Ontology) of aging-associated and aging-independent processes identified by GSEA analysis.

### Apelin activity affects the development of the HPT axis

Given that our GO and KEGG enrichment analyses pointed to reproduction and hormonal regulation, we next sought to directly test this hypothesis by examining genes central to the hypothalamic-pituitary-thyroid (HPT) axis regulation and function. We analyzed the expression of genes involved in growth and reproduction, including growth hormone (*gh*), luteinizing hormone (*lh*), follicle-stimulating hormone (*fsh*), prolactin (*prl*) and proopiomelanocortin (*pomca*), as well as genes involved in thyroid hormone regulation, including thyrotropin-releasing hormone (*trh*), thyroid-stimulating hormone (*tsh*), and thyroglobulin (*tg*). We first investigated the change in gene expression in the *apelin* mutant, severe phenotypes, at 5 dpf by *in situ* hybridization. While *gh* and *prl* expression are largely unaffected in mutant condition (Fig. 3A), we could not detect *lh* and *fsh* expression in the pituitary or hypothalamic *trh* expression in any of the experimental conditions with regular *in situ* hybridization protocol. However, we observed that hypothalamic *pomca* expression is largely reduced in the mutant (Fig. 3A), correlating with dysregulation of stress and reproductive processes observed in the mutant. In addition, we identified that *tg* expression is affected in the mutant (Fig. 3A), suggesting that thyroid signaling may be dysfunctional. Supporting the above observation and a role for Apelin in the thyroid system development, we demonstrated that *tg* expression is strongly downregulated in the Apelin gain-of-function (10 µM of 4-OHT from 3 to 5 dpf) condition (Fig. 3A). We next performed hybridization chain reaction (HCR) fluorescent *in situ* to quantify gene expression in loss and gain of *apelin* function. Of note, HCR *in situ* being more sensitive, we detected *trh* and *lh* expression (Fig. 3B, C), but still not *fsh* mRNA in the pituitary. Following confocal imaging and 3D reconstruction and segmentation of *trh*, *tsh* and *tg* expression, we demonstrated that *trh* cell number is significantly reduced in the loss (severe phenotype) and gain of function experiment, while *tsh* reduction is significantly observed only in the gof (Fig. 3B). Interestingly, the most significant difference was observed using *tg* as a marker of the thyroid gland. We quantified a reduction in thyroid size (volume after 3D reconstruction) in both the loss and gain-of-function conditions (Fig. 3B). Strikingly, while the thyroid morphology is different in loss and gain of Apelin function (Fig. 3B), both phenotypes converge on a reduction of the size, underscoring whether the proper level and timing of *apelin* expression is critical for the normal development of the thyroid organ. We next used fluorescent *in situ* hybridization to analyze *lh* and *pomca* gene expression (Fig. 3C). We identified that the number of *lh+* cells in the pituitary and *pomca* expression are affected by Apelin activity (Fig. 3C). Importantly, our data indicates that 4-OHT treatment impacts significantly the expression of *pomca* in both the pituitary and the hypothalamus (Fig. 3C). This observation implies that the reduction of the *pomca*+ pituitary volume is independent of *apelin* GOF, but rather a result of the 4-OHT treatment. Finally, similarly to the thyroid organ, both loss and gain of *apelin* function have a similar impact on the formation of *lh*+ gonadotropic cells in the pituitary gland. Together, this analysis demonstrates that the Apelin peptide controls the homeostasis of the HPT axis, in particular the expression of neuropeptides regulating pituitary and thyroid-dependent signaling, two structures known to be essential for growth, fertility, and aging.

**Figure 3.**
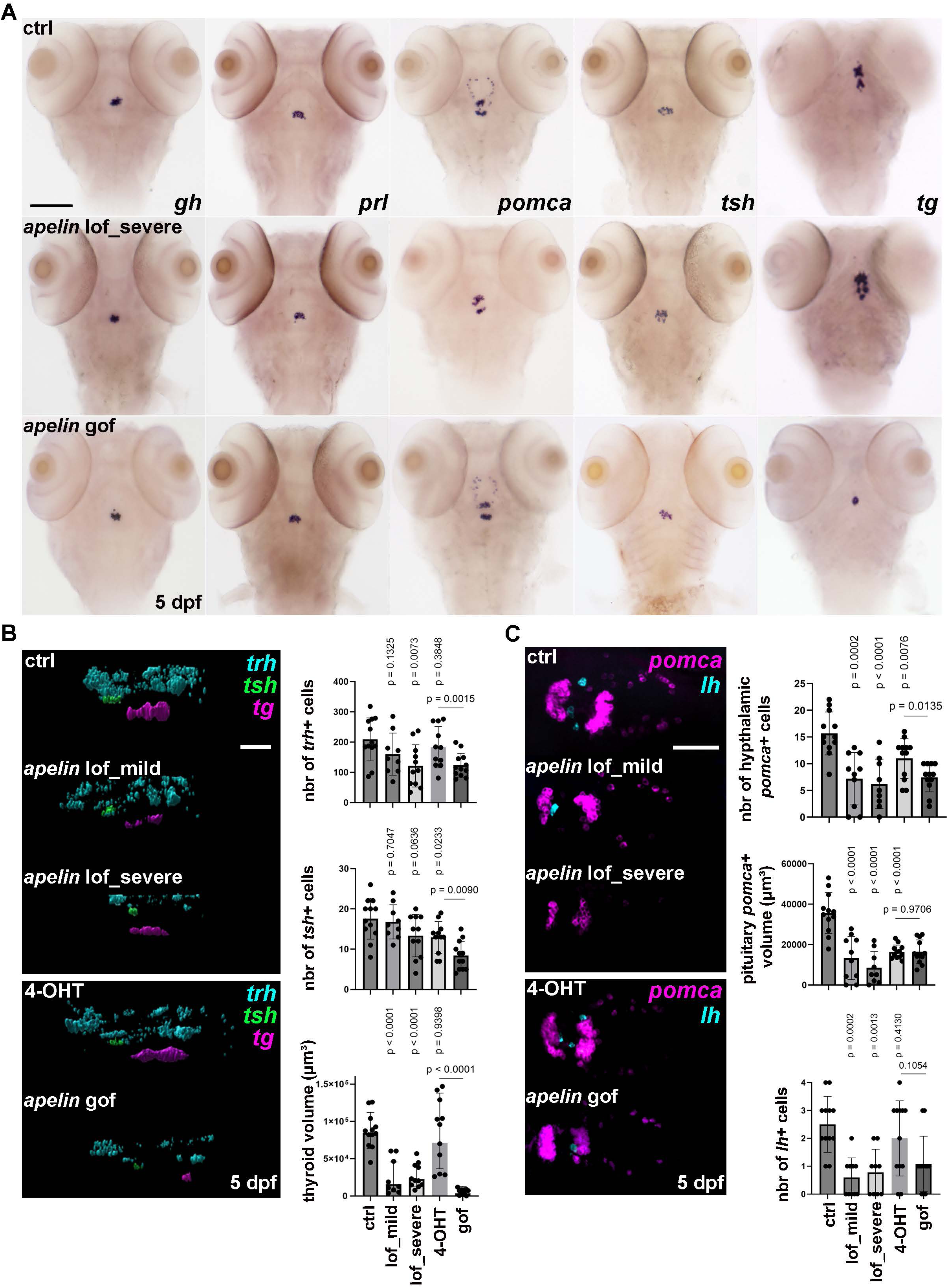
Apelin dysregulation affects hypothalamo-pitituary-thyroid axis development. **(A)** Gene expression analysis of *gh, prl, pomca, tsh,* and *tg* by ISH from top to bottom, control, *apelin* loss-of-function severe phenotype, and *apelin* gain-of-function at 5 dpf. Control and *apelin* overexpression have been treated with 4-OHT from 3 to 5 dpf. **(B)** Analysis of *trh, tsh, tg HCR in situ* hybridization at 5 dpf. Control and *apelin* overexpression have been treated with 4-OHT from 3 to 5 dpf. Quantification of *trh+ cells, tsh+ cells* number and thyroid volume *tg+* in wild-type (n= 12), *apelin* mutant mild (n= 9) and severe phenotype (n= 11), wild-type treated 4-OHT (n= 11) and *apelin* gain-of-function (n= 12). **(C)** Analysis of *lh, pomca HCR in situ* hybridization at 5 dpf. Control and *apelin* overexpression have been treated with 4-OHT from 3 to 5 dpf. Quantification of *lh+* cells in the pituitary gland, hypothalamic *pomca+* cells, and pitituary volume *pomca+* in wild-type (n=12), *apelin* mutant mild (n=10) and severe phenotype (n= 9), wild-type treated 4-OHT (n= 12) and *apelin* gain-of-function (n=12). Significance is determined by two-sided unpaired Student’s T-test and Mann-Whitest non-parametric test. Error bars represent s.d. Scale bars: 100 μm.

### *atoh8* is a downstream target of the Apelin peptide regulating HPT axis development

To identify genes regulated by Apelin signaling at the larval stage, we performed bulk RNA-seq on larvae (25 larvae/sample, 4 replicates) in the loss of *apelin* function (Fig. 4A and Fig. S3A, B). To further assess Apelin’s biological function, we also included a gain-of-function condition in this analysis (Fig. 4B). We observed similar patterns of gene dysregulation in opposite direction when we compared the *apelin* KO and upregulation conditions (Table S21-24). In particular, but not limited to, *atoh8*, *klf9*, *fkpb5*, *vegfaa*, *vegfba* and *mpeg1.1* behave as transcriptional targets of Apelin signaling with LOF and GOF in larvae that were also similarly dysregulated independently of aging in the different adult *apelin* mutant tissues (Table S16-18 and S21-24). Interestingly, we also identified *trhde* and *tg*, that are both involved in thyroid signaling and exhibit opposing pattern of dysregulation in the loss and gain-of-function conditions. Among all these genes, *atoh8* has been of particular interest. Atoh8 is a bHLH transcription factor consistently downregulated across all three tissues in an aging independent manner and represents a non-hormonal and non-metabolic factor affected by Apelin signaling. Atoh8 is required for normal retinal lamination, myogenic patterning and angiogenesis in zebrafish, whose morphant knockdown produces reduced eye size, disorganized somite/muscle architecture and cardiovascular defects ^79,80^; biological processes that are also affected in the zebrafish larvae mutant for the *apelin* gene. This observation suggests that Atoh8 may be a transcriptional effector, downstream of the Apelin signaling pathway. To test this hypothesis, we injected the *apelin* mRNA in zebrafish embryos and observed that it leads to upregulation of the *atoh8* gene at 12 hpf, indicating that *atoh8* is a target of the apelinergic system (Fig. S3C). Next, we attempted to alleviate the *apelin* mutant phenotype with *atoh8* mRNA expression to assess whether *atoh8* regulation by Apelin participates in the mutant phenotype. We first determined that contrary to *apelin* LOF, *atoh8* overexpression has little effect on early development (Fig. S3D-F) and observed that *atoh8* expression in the *apelin* mutant genetic background can attenuate the embryonic development defects (Fig. S3G, H). We next quantified the *apelin* mutant phenotype rescues by *atoh8* mRNA and observed significant shift in the proportions of the mild phenotype (42.8% vs 12.99% in heterozygous incross vs rescue respectively) phenotype toward a normal phenotype (35.7.8% vs 67.8% in heterozygous incross vs rescue respectively) for both heterozygous (44% vs 51.28%) and homozygous (13% vs 24.54% respectively) mutants (Fig. S3D). Interestingly, this transition is accompanied by a ∼1.8-fold reduction of the *apln*+/+ genotype in normal phenotype category, supporting a positive effect of *atoh8* expression to alleviate phenotypes associated with the *apelin* mutation (Fig. S3D). We next asked whether *atoh8* expression in the *apelin* homozygous mutant harboring a normal phenotype after rescue could also reverse some of the HPT defects observed in the *apelin* homozygous mutants. Through 3D reconstruction, segmentation and quantification at 3 dpf, we demonstrated that *atoh8* mRNA expression can significantly increase *trh*+ and *tsh*+ cell number as well the volume of the thyroid in the mutant (Fig. 4C, E), indicating not only *atoh8* alleviates gross morphological defects, but also partially rescue the development of the HPT axis. Similarly, we observed a positive impact of *atoh8* mRNA expression on *pomca*+ expression in the hypothalamus of homozygous *apelin* mutant while we determined a non-significant effect on the pituitary volume and the number of *lh*+ cells in the pituitary at this stage (Fig. 4D, F). The above observations suggest that Atoh8 transcription factor may act on biological processes, such has the HPT axis, that are also downstream of Apelin signaling. Because studies of the role of *atoh8* during early development using knockdown and loss of function are still controversial ^79,80^, we wanted to confirm the above hypothesis. We generated *atoh8* crispant larvae (Fig. S3I-M) and identified phenotypes similar to those reported previously in the *atoh8* morphant ^79,81^, supporting a potential role for the Atoh8 transcription factor downstream of the Apelin signaling pathway (Fig. S3G). We next quantified the role of Atoh8 in the development of the HPT axis using gain and loss of function experiments. As mentioned above, overexpression of *atoh8* mRNA has little impact on early development with no obvious gross defects (Fig. S3D, F). However, we observed a mild but significant increase in the number of *tsh*+ cells and *tg*+ thyroid volume, and a reduction of *pomca+* pituitary gland volume after *atoh8* expression (Fig. 4C-F). In contrast, *atoh8* crispants harboring a normal or severe phenotype at 3 dpf have a reduced *tg*+ thyroid volume but show an increase of *pomca*+ volume of the pituitary gland (Fig. 4C-F), indicating that the proper formation of this neurendocrine axis requires *atoh8* function. In addition, the number of *tsh*+ and *lh*+ cells in the pituitary of *atoh8* crispants is only slightly, but significantly, affected (Fig. 4C-F). Finally, the number of *trh*+ and hypothalamic *pomca*+ cells are only reduced in the *atoh8* crispants showing a severe phenotype, suggesting that the hypothalamus is less sensitive to the loss of *atoh8* function compared the pituitary and thyroid glands (Fig. 4C-F). Taken together, these findings suggest that Apelin signaling affects development and function of the major hypothalamic-pituitary axes, at least partially through its Atoh8 downstream effector, potentially regulating a variety of biological processes associated with tissue homeostasis and aging including, but not limited to, the thyroid (HPT) axis governing metabolism, the adrenal/interrenal (HPA/I) axis regulating stress response, and the gonadal (HPG) axis controlling reproduction.

**Figure 4.**
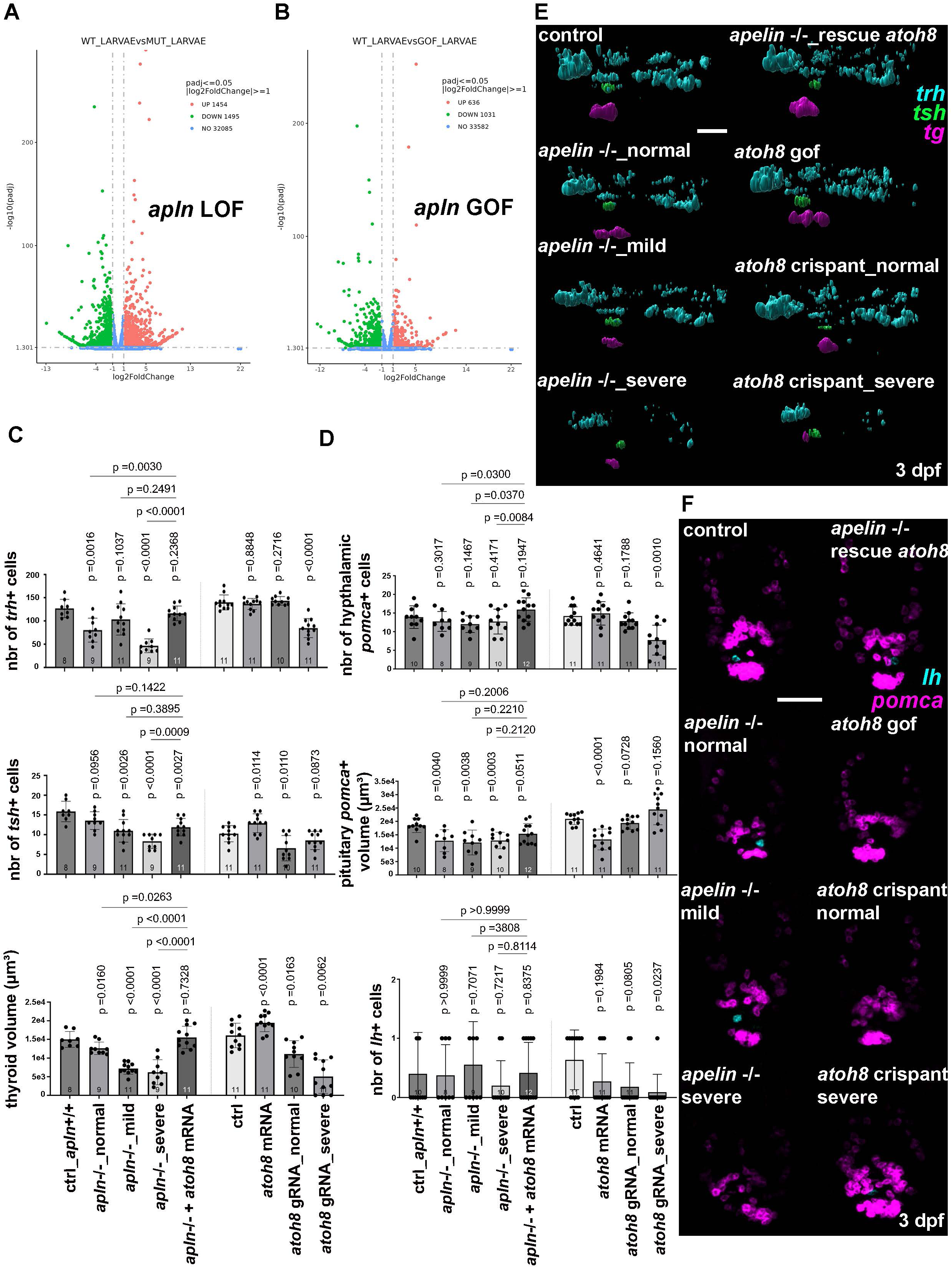
*atoh8* is a downstream regulator of Apelin signaling that partially rescues the *apelin* mutant phenotype. **(A, B)** Volcano plots of downregulated (red) and upregulated (green) genes identified by GSEA in *apln* loss-of-function (A) and *apln* gain-of-function (B) larvae at 6 dpf. **(C, E)** Quantification (C) and representative 3D reconstructions (E) of *trh, tsh* and *tg* HCR *in situ* hybridization at 3 dpf, showing the number of *trh+* and *tsh+* cells and thyroid volume (*tg+*), in control *apln +/+* (n= 8), *apln* -/-normal (n=9), mild (n=11) and severe (n=9) phenotypes, *apln* -/- + *atoh8* mRNA injection (n=11), control uninjected (n=11), control injected with *atoh8* mRNA (n=11), and *atoh8* crispants normal (n=10) and severe phenotype (n= 11). **(D, F)** Quantification (D) and representative 3D reconstructions (F) of *lh* and *pomca* HCR *in situ* hybridization at 3 dpf, showing the number of *lh*+ cells in the pituitary gland, hypothalamic *pomca*+ cells and pituitary volume (*pomca+*), in control *apln +/+* (n=10), *apln* -/- normal (n=8), mild (n=9) and severe (n=10) phenotypes, *apln* -/- injected with *atoh8* mRNA (n=12), control uninjected (n=11), control injected with *atoh8* mRNA (n=11), and *atoh8* crispants normal (n=11) and severe phenotype (n= 11). Statistical significance is determined by two-tailed unpaired Student’s t-test and Mann-Whitney test, non-parametric. Error bars represent s.d. Scale bars: 100 µm.

## DISCUSSION

Our study reveals an underappreciated role for Apelin signaling for the normal development of the hypothalamic–pituitary–thyroid (HPT) axis in zebrafish and establish Atoh8 as a transcriptional effector downstream of the Apelin pathway whose restoration partially rescues the mutant phenotype. The genetics of this functional requirement is very sensitive to Apelin dosage during early embryogenesis, where reducing *apelin* expression in heterozygous is sufficient to produce a phenotype mimicking the homozygous loss of function. Yet removing it entirely neither worsens the defects nor makes them fully penetrant, so heterozygous and homozygous mutants develop the same spectrum. Incomplete penetrance in homozygotes embryos implies that apelin loss is buffered by other molecular components: either maternally deposited transcript or peptide from heterozygous parents, or residual activation of AplnR by Elabela during early development. Similar sensitivity to gene dosage is not unusual among secreted regulators of vascular morphogenesis, as heterozygosity for a *Vegfa* null allele is lethal at embryonic day 9.5 in mouse, and heterozygous deletion of *Dll*4 produces prominent though variable arterial defects ^100^, leading Gale et al. to conclude that vascular development is as sensitive to subtle changes in Dll4 dosage as it is to *Vegfa* dosage. A plausible mechanistic basis is that apelin is known to function as a positional cue, where a ligand instructs directional cellular behavior through a spatial gradient. This interpretation is supported by work on sprouting angiogenesis ^102^, in which vascular tip cells elongate and migrate directionally toward and along Apelin-expressing neural progenitors, and in which restoring neural, but not vascular, *apelin* expression is sufficient to correct the angiogenic defects of *apelin* mutants.

We identified a previously unreported phenotype on the HPT axis in the zebrafish *apelin* mutant. That Apelin acts in this region is consistent with its documented distribution in the mammalian neuroendocrine hypothalamus ^103,105^, where it alters pituitary hormone output in several species, raising ACTH and Corticosterone levels while lowering Prolactin, LH and FSH in rat ^104^. In teleosts, by contrast, the hypothalamic apelinergic system has been studied almost exclusively in the context of energy balance: Apelin is an appetite-regulating peptide expressed in the fish brain ^106^, hypothalamic Apelin falls after feeding and rises with fasting ^109^. While a developmental requirement for Apelin signaling in the teleost HPT axis has not, to our knowledge, been reported, precedents for hypothalamic peptides acting on the zebrafish hypophysis does exist, since MCH, POMC, AGRP, NPY and Oxytocin neurons target the pituitary gland and/or the neurohypophysis blood vessel portal to control complex behaviors such as background adaptation, stress response, feeding and reproduction ^107,108,110–112^.

The pituitary is a direct modulator of the thyroid gland activity, and both organs share similar functions regulating complex behaviors mentioned above. However, the vasculature development and organization differ between these two endocrine glands. The pituitary’s vessels are specialized for neurovascular/endocrine communication, whereas thyroid vessels are arranged around follicular units to support thyroid hormone production and secretion. While the pituitary develops before the establishment of its specialized neurovascular network, with vascularization occurring progressively during pituitary development ^108^, the zebrafish thyroid primordium develops as a follicular endocrine organ that follows the blood vessels dispersing along the pharyngeal midline ^113^. These developmental differences between the endocrine-vascular interactions of the two glands raised different hypotheses, that are not mutually exclusive, regarding the role of Apelin signaling in these two processes of morphogenesis. Whether Apelin can act in a cell autonomous manner on the endocrine gland to control its development, or it can indirectly affect their formations by modulating morphogenetic cues from adjacent tissues. Because of the well-established role of Apelin in blood vessel development ^14^ and because of the close endocrine-vascular association ^108,113^, the vasculature is a prime candidate supporting the latter hypothesis. Future studies using *apelin* cell specific knock-out, rescue or gain of function would be required to address autonomous versus non-autonomous developmental mechanisms underlying the HPT phenotype observed after perturbation of Apelin expression.

Our work also supports a role for Apelin-dependent control of blood vessel development via regulation of Vegf-A expression, as *vegfaa* is downregulated in the different tissues analyzed with loss of *apelin* function. Interestingly, *vegfba* follows an opposite trend of expression being consistently overexpressed in the *apelin mutant*. While we are cautious about the interpretation of these results, it may represent a compensatory response of an orthologue gene to the altered expression of the other. This mechanism has been previously documented for *Apelin* loss of function leading to de-repression of other VEGF family members in lymphatic endothelium, where *Apelin* deficiency raises VEGF-C and VEGF-D expression ^114^. Another possible explanation regarding this phenotype is that *vegfaa* and *vegfba* orthologues share a different function. VEGF-B has been reported to be responsive to stress and challenges, acting as a cell survival and anti-apoptotic factor rather than a major pro-angiogenesis molecule ^117^. This observation correlates with our RNA-seq results indicting that stress response is elevated, in the absence of the *apelin* gene, as indicated by overexpression of genes involved in the stress response pathway such as *kfl9* and *fkbp5* ^83^. Apelin signaling being a major player in cardiovascular development and aging, a simple hypothesis would be that defects leading to premature aging, and potentially hypoxia, will increase cellular stress in the *apelin* mutant. However, because Apelin has pleiotropic roles during development and aging, it is likely that the high stress in the mutant represents a more complex response driven by different tissues and organs.

APLNR/APJ expression within components of the HPG and HPT axis supports a role for apelinergic signaling in the development and functional regulation of neuroendocrine homeostasis ^1,25^. APLNR is expressed in discrete hypothalamic and pituitary regions involved in neuroendocrine regulation, including areas associated with TRH neurons ^115,116^. APLNR expression has also been detected in pituitary and thyroid tissues ^25,118^, raising the possibility of local apelinergic regulation at multiple levels of the HPT axis. However, although these anatomical associations support a modulatory role for APLNR in HPT-axis maturation and function, direct evidence linking APLNR signaling to HPT development remains limited. Our study provides such functional evidence linking Apelin expression to developmental defects of the HPT axis, positioning Apelin as a metabolic and neuroendocrine regulator that contribute to the TRH–TSH–thyroid hormone pathway.

While the role of Apelin in the development and function of the HPT axis is potentially indirect and non-cell autonomous, we identified a downstream transcriptional effector of the Apelin signaling pathway that has the potential to act in a cell autonomous manner. Atoh8 is a bHLH transcription factor that regulates gene expression positively or negatively in a context-dependent manner during development and cellular homeostasis ^119^. Through its conserved bHLH domain, Atoh8 binds E-box DNA elements and interacts with other transcription factors and signaling effectors ^120^. Similarly to Apelin, Atoh8 has been implicated in the regulation of diverse biological processes, including neuronal and retinal differentiation, pancreatic endocrine development, skeletal myogenesis, endothelial differentiation, placental trophoblast specification, and renal development ^79,121–123,125^. Notably, its effects can vary substantially among tissues and disease states, reflecting differences in transcriptional partners, signaling context, and Atoh8 isoform expression ^119,124,127^. Thus, Atoh8 is increasingly recognized as a multifaceted transcriptional regulator that integrates developmental and extracellular signals to modulate cellular differentiation, metabolism, and stress responses.

Atoh8 has an important role in skeletal, muscular and vascular development ^79,129^, In human endothelial cells, ATOH8 is induced downstream of BMP-9/SMAD1 signaling ^129^ and contributes to endothelial differentiation, proliferation, migration, and vascular tube formation ^126^. In vivo, loss of *ATOH8* in mice results in delayed retinal vascular outgrowth and a pulmonary vascular phenotype resembling pulmonary arterial hypertension, supporting a role for ATOH8 in vascular maturation and adaptation to hypoxic stress. Notably, ATOH8 interacts functionally with HIF-2α and appears to attenuate hypoxia-responsive transcription in the pulmonary circulation, linking BMP signaling, oxygen sensing, and vascular homeostasis ^129^. In zebrafish, in addition to its role during cardiovascular angiogenesis ^81,126^, Atoh8 is involved in skeletal and muscle development, phenotypes correlating with it expression pattern during embryonic and larval development ^79,80^. However, the exact role of Atoh8 in these biological processes is still debated in the field as contradictory reports have been published ^81,128,130^, in particular regarding its function during cardiac development. In our experiments, *atoh8* crispants harbor phenotypes reminiscent to the morphant phenotypes, but not identical to those in stable established *atoh8* knock-out mice and zebrafish ^80,81,128,130^. These discrepancies may lie in the different genetic compensatory response between transient loss of function established with morphants and crispants compared to full stable knock-out mutants. This phenomenon has been reported extensively in fish and mice, where stable loss of function in the germline can trigger a genetic compensatory response masking the phenotype ^131–134^.

ATOH8 is functionally connected to the Apelin signaling pathway through its role in endothelial and vascular biology ^135–137^. Importantly, ATOH8 is positioned within the BMP9/BMP10–ALK1–SMAD1/5/8 signaling network ^129^, a pathway that regulates vascular homeostasis. Thus, ATOH8 may participate in an interconnected regulatory network in which BMP–SMAD signaling, ATOH8, hypoxia-responsive transcription, and the APLN/APLNR axis converge to coordinate angiogenic responses. The co-expression of ATOH8 and APLNR in endothelial cell populations ^138^ further supports the biological proximity of these pathways. The above information is supported by our transcriptomics analysis and functional experiments, although the exact regulatory mechanism of *atoh8* downstream of Apelin signaling to be identified. While a direct requirement for Atoh8 in development of the HPT axis has not been previously established, our work suggest that gain of *atoh8* function can partially rescue the *apelin* mutant phenotype. Whether this effect is direct or indirect in the neural and endocrine tissues still need to be determined. However, because *atoh8* is expressed in multiple embryonic tissues and regulates differentiation, BMP/TGF-β signaling and vascular development, it may represent a candidate regulator of the HPT-axis organogenesis. Its specific roles in hypothalamic, pituitary, or thyroid development and in subsequent neuroendocrine signaling remain to be defined experimentally.

In conclusion, we showed that *apelin* expression level must be tightly controlled within a narrow range for normal zebrafish development, as both loss and gain of function lead to similar phenotypic defects. Some of these phenotypes persist into adulthood: both *apelin* mutant and overexpressing fish are largely infertile and die early, with homozygous mutants reaching 50% mortality by 180 dpf. Transcriptomic profiling of adult brain, eye, and trunk distinguishes an aging-associated gene set enriched for hormone transport and secretion from an aging-independent set enriched for sexual development, and places 12-month-old mutants closer to 48-month-old wild-type fish than to their age-matched siblings. At the tissue level, both loss and gain of function reduced *trh*+ cell number and thyroid volume, identifying the hypothalamic-pituitary-thyroid axis as a target of Apelin signaling. We identified *atoh8* as an effector in this process: it is induced by *apelin*, downregulated in the mutant, and its re-expression in *apelin* lof partially restores *trh*+ and *tsh*+ cell numbers as well as thyroid volume. Together, these findings identify Apelin signaling and its effector *atoh8* as regulators of hypothalamic-pituitary-thyroid axis development and function, and suggest that their dysregulation may underly reproduction deficits and premature aging.

## MATERIALS AND METHODS

### Study design and statistical analysis

Research subjects (larval stage and genotype), number and treatments applied are defined in the figures and/or legends. Experimental analyses were performed with a minimum of 3 biological replicates (and a minimum of 3 technical replicates when applicable), each zebrafish adult sample or larvae is considered as a biological replicate. Effect sizes for each comparison were calculated as Hedges’ g, as recommended for groups with fewer than 20 observations ^139^. Effect sizes were interpreted following Cohen’s ^140^ conventions, with |g| ≥ 0.8 considered large. The number of biological replicates and larvae used for each specific experiment as well as the statistical test used to calculate significance is indicated in the figure legends. For statistical analysis, p-values are indicated in the figures. Comparisons were conducted to determine significant differences between control and experimental conditions. Statistical analyses were performed using GraphPad Prism 11. The normality of data distribution was assessed using the Shapiro-Wilk test and visualized with quantile-quantile (Q-Q) plots. Homogeneity of variance between groups was evaluated using F-tests. Based on these analyses, continuous data were compared using Student’s t-test, Welch’s t-test, or the Mann-Whitney test, and categorical data were compared using Fisher’s exact test, as indicated in the figure legends. Statistical significance was set at p < 0.05, and exact p-values are shown in the figures. Statistical significance was set at p < 0.05, and exact p-values are shown in the figures.

### Zebrafish lines and developmental conditions

Experiments were performed in accordance with animal care guidelines. Our IACUC animal protocol (AUP#22-06) is approved by the MDI Biological Laboratory. For imaging analysis, larvae were fixed overnight at 4°C in 4% paraformaldehyde/1xPBS, after which they were dehydrated through an ethanol series and stored at −20°C until use. The generation and validation of Tg(ubi:Lox-stop-Lox-*apelin*) ^82^ and Tg(*503unc:creERT2*) ^83^ have been reported previously.

### CRISPR/Cas-9 generation of a zebrafish *apelin* and *atoh8* mutant

To generate and *atoh8* mutants, we took advantage of a previously described genome-editing method ^141^. Specifically, we used tracrRNA and crRNA (IDT) to form functional gRNA duplexes targeting the ORF (*apln*: 5’-GAATGTGAAGATCTTGACGC-3’; *atoh8*: 5’-TGTCTCCTGACACCGTGTTC-3’ and 3’-GTTGACAGTCTTTTCGGAGA-5’) Co-injection of the gRNAs with the Cas-9 protein (IDT) resulted in indels in the *apln* and *atoh8* genes coding sequences, identified by Sanger sequencing and analyzed with ICE software (Synthego) in F0 injected embryos. F0 carriers for mutations in the *apelin* locus in the germline were identified by sequencing the alleles on the clutches and were then outcrossed to obtain F1 heterozygous mutants. Mutations (indels) leading to a premature stop codon were identified by ICE analysis (Synthego) of the Sanger-sequencing results. All subsequent F# *apelin* heterozygous and homozygous mutants were confirmed by PCR genotyping with *apln-*crispr-ID forward: “5’-CTGTCACACACTTGCACAC-3” and reverse: “5’-GCACAATAACAAGCCAAACCTC-3’” primers followed by sanger sequencing. The *apelin* mutant harbor a 26-bp insertion leading to a stop codon at position 18 of the peptide as detailed in our previous study ^82^.

### mRNA synthesis and microinjection

Capped *apln* or *atoh8* mRNA were synthesized from NotI-linearized pCS2+ vector containing the coding sequence using the mMESSAGE mMACHINE SP6 Transcription Kit (Invitrogen) and then purified on column (Qiagen). RNA integrity was confirmed by agarose gel electrophoresis and quality and concentration was measured by NanoDrop. For microinjection, mRNA was diluted in nuclease-free water containing 0.1% phenol red to a final concentration of 150 ng/µl.

### Pharmacological treatments

Stock solution of 1-phenyl 2-thiourea (PTU; Sigma) was prepared at 25X (0.075%). PTU 1X working solution was used to inhibit pigmentation in zebrafish embryos. A stock solution of hydroxytamoxifen (4-OHT; Sigma) and Tamoxifen (Sigma) were prepared at 10 mM in 100% DMSO (Sigma). For induction of LoxP-site recombination, zebrafish larvae were treated with 4-OHT at a concentration of 10 µM for a duration of 48h and zebrafish adults were treated with Tamoxifen at a concentration of 10 µM for a duration of 24h weekly.

### Gene-expression analysis

*In situ* hybridizations were performed as previously described ^142^. The ORFs of *gh, prl, pomca, tsh, and tg,* were cloned in a pCS2+ vector using zebrafish cDNA. Antisense DIG-labelled probes were transcribed using T3 polymerase (Promega) and the linearized pCS2+ plasmid containing the ORF as a template. DIG probes were detected using anti-DIG antibody (Roche), and *in situs* were revealed using BCIP reagent (Roche). HCR fluorescent *in situ* experiments were performed using probes, fluorophores, and protocols provided by Molecular Instruments (HCR™ RNA-FISH (v3.0)).

Total RNA samples for bulk RNAseq analysis were performed using biological replicates to ensure reproducibility (4 biological replicates (2 males and 2 females) for the adult analysis and 3 biological replicates ∼25 larva/sample for the larval analysis). Trizol (Invitrogen) was used to homogenize cellular extracts with the TissueLyzer (Qiagen). RNA purification was performed as described above. For bulk RNAseq analysis, library preparation and sequencing were carried out by NovoGene using the Illumina NGS platform. Indexing and alignment were performed using HISAT2 (v2.0.5) ^143^ against the GRCz11 (*Danio rerio*) reference genome obtained from Ensembl. Mapped reads were assembled using StringTie (v1.3.3b) ^146^, and quantification was performed using FeatureCounts (v1.5.0-p3) ^144^. Differential expression analysis was conducted in the R environment using DeSeq2 ^145^, and Gene Set Enrichment Analysis (GSEA) was performed using clusterProfiler ^147^. All visualizations were generated using ggplot2.

The gene lists were then intersected using InteractiVenn ^148^ and orthology assignments were obtained with g:Profiler ^149^, and gene set enrichment analysis was performed with ShinyGO ^150^. Given the limited number of genes in the input lists, a relaxed significance threshold (p > 0.1) was applied for the enrichment analysis. The resulting enrichment terms are reported as descriptive only. RNAseq data are available via the GEO database and the accession number GSE351298 and GSE351299.

### Locomotion (larvae) and swimming (adults) analysis

Larvae were treated with hydroxytamoxifen or DMSO at 10 μM from 5 dpf to 7 dpf followed by multiple washes with fresh water to remove chemicals. Individual larvae were placed in 48-well plates containing 500 μL of medium. Locomotion assays were performed using Zebrabox® and recorded using Viewpoint® quantization software. Parameters used were: threshold 19, freezing 5, and burst 6, with data integration every 2 hours. A backlight (20%) and toplight (3%) were activated during the assay. Total locomotive activity per 2-hour period was calculated by summing small and burst movements and comparing them with freezing activity. Maximum values did not exceed 7,200 seconds (2 hours) per integration point. Raw data were transformed into percentage of activity and analyzed using GraphPad Prism. Multiple unpaired t-tests with the False Discovery Rate (FDR) method were performed, analyzing mean and variance for each population row. Each data point represents one individual. For visibility the graph is represented with the mean of the population per condition (16 larvae/condition).

Swimming tunnel preparation, calibration, and maintenance were adapted from *Burris et al* as reported previously ^83^. The swim performance assay was modified to use velocity increments of 5 cm/s every 3 minutes for a total duration of 60min, with fish size positioned at 3 cm in length and an average of 5 zebrafish swimming simultaneously in the swimming chamber. This group size was selected to avoid conflict between zebrafish while enabling accurate recording of swimming performance for each individual fish. Before the performance assay started, zebrafish were acclimated for 15min in the swimming chamber with a velocity of 30cm/s.

### Beating heart analysis

Samples were anesthetized with tricaine and positioned in a glass container with a glass bottom. Images were acquired using a Zeiss Axiozoom v16 unit, equipped with an objective magnification 1x, plan Z, and numerical aperture (NA) of 0.25, immersion medium Air (dry) and zoomed 160x. Each fish was recorded for 1min then place back in the incubator with their siblings. Images were collected every 10ms, using an exposure time of 8ms for a total duration of 1 minute. To count heartbeats, the Zplot axis was used and peaks were counted manually.

### Body length analysis

Larvae were fixed overnight at 4°C in 4% paraformaldehyde/1xPBS, after which they were rehydrated through an ethanol series and submerge in 70% glycerol solution then imaged using a Zeiss Axiozoom v16 unit, equipped with an objective magnification 1x, plan Z, and NA of 0.25, immersion medium Air (dry) and zoomed 50x.

Adults were anesthetized with tricaine then placed on a white sheet with a ruler beneath it and taken in picture for future assessment. Length was obtained by drawing a straight line from the tip of the fin and head down to the ruler then.

### Image acquisition, 3D reconstruction and image analysis

Images were acquired using a point scanning confocal unit (LSM 980, Carl Zeiss Microscopy, Germany) on a Zeiss Axio Examiner Z1 upright microscope stand (409000-9752-000, Carl Zeiss Microscopy, Germany) followed by 3D reconstruction using Imaris software. Volume was reconstructed by creating a mask of fluorophore-positive tissue and was adjusted manually to correct for auto fluorescent structures such as blood vessel or blood cells.

Volumes thyroid (*tg*), Pituitary gland (*pomca+*), and number of cells (*trh*) at 3 or 5 dpf, were extracted from the ‘Statistics’ module and uploaded to GraphPad Prism for statistical analysis. Condensed cluster of cells were counted as one cell. Final 3D reconstruction images were extracted from Imaris software using Snapshot module and were exported as .PNG files.

Counting for *lh*+ and *tsh+* in the pituitary gland and *pomca+* cells in the hypothalamus were manually extracted from 2D visualization with the 2D view from Imaris software. Counting was replicated three times and averaged to provide final count then uploaded to GraphPad Prism for statistical analysis.

### Image creation

Created in BioRender. Menard, R. (2027) https://BioRender.com/g7rvcqa

Created in BioRender. Menard, R. (2027) https://BioRender.com/v18ced5

## Supporting information

SuppFigures

## ACKNOWLEDGMENTS

This study was supported by Institutional Development Awards (IDeA) from the National Institute of General Medical Sciences of the National Institutes of Health under grant numbers P20GM103423 (MDIBL), P20GM104318 (MDIBL), and P20GM144265 (RM). Funding was also provided by the Morris Discovery Fund (RM).

Image collection, processing, and analysis for this manuscript was performed with the assistance of Dr. Frederic Bonnet and the MDI Biological Laboratory’s Light Microscopy Facility (RRID:SCR_019166), which is supported by an Institutional Development Award (IDeA) from the National Institute of General Medical Sciences of the National Institutes of Health under grant number P20GM103423.

## Authors’ contributions

*Conceptualization*: Romain Menard and Romain Madelaine. *Investigation*: Romain Menard, Anindita Neog, Caroline Halluin, Federic Bonnet and Romain Madelaine. *Data curation*: Romain Menard and Romain Madelaine. *Formal analysis*: Romain Menard and Romain Madelaine. *Funding acquisition*: Romain Madelaine. *Supervision*: Romain Madelaine. *Writing – original draft:* Romain Menard. *Writing – Review & Editing:* Romain Menard, Romain Madelaine.

## Data availability

All data, reagents, and genetic tools presented in this manuscript are available to the scientific community.

All the authors have no competing interests.

---

**Figure S1. *apelin* mutants and overexpression display reproductive, locomotive and aging phenotypes. (A)** Survival analysis of *apelin* +/+ (n=211) and *apelin* +/- (n=340) larvae at 5 dpf (B) Quantification representing the number of crossings (displayed on the graph) between wild-type, *apelin* mutant homozygous, heterozygous and overexpression fish, males or females. **(C)** Percentage of fertilized (n=3) vs unfertilized (n=573) eggs. **(D)** Characterization of the phenotypes and genotypes followed from 3 to 5 dpf of wild-type (top left), *apelin* mutant (bottom left), wild-type treated with 4-OHT 10 µM (top right), *apelin* gain-of-function (bottom right) clutches. Each plot is read from the center outward. The center ring shows the distribution of phenotypes at 3 dpf, with percentages summing to 100% of the sampled population. Each 3 dpf phenotype group was then followed individually and re-sorted at 4 dpf (middle ring), and each resulting 4 dpf subgroup was followed and re-sorted at 5 dpf (outer ring). Each segment therefore sits directly outside the group it came from. Percentages in the 4 and 5 dpf rings are relative to the parent segment. For example, in the *apelin* mutant plot, 53.82% of larvae were mild at 3 dpf. At 4 dpf, this group split into mild (93.41%) and severe (6.59%). The 4 dpf mild subgroup then split at 5 dpf into normal (10.53%), mild (83.63%), severe (2.92%), and dead (2.92%). Phenotypes and genotypes of single larva can be visualized as follows: dead (red), severe (orange), mild (yellow), normal (green), wild-type (light grey), heterozygous (medium grey) and homozygous (dark grey). Inner ring: global genotype distribution (%) across the sampled population. Outer ring: genotype frequencies (%) within each phenotype subpopulation. **(E)** Quantification of heartbeat of WT (n=12), apelin overexpression mild (n=12) and severe (n=12) phenotypes treated with 4-OHT 10 µM and apelin mutant mild (n=12) and severe (n=12) phenotypes. Statistical significance is determined by Mann-Whitney test, non-parametric. **(F)** Quantification of locomotion activity in zebrafish larvae. Wild-type (n=16) and *apelin* overexpression (n=16) larvae treated with 4-OHT, and *apelin* mutant screened on mild phenotypes (n=16). Larvae were treated with 4-OHT for 48 hours between 5 and 7 dpf during the test. Statistical significance is determined by multiple t-test comparison with False Discovery Rate and individual variance for each time point, two-tailed, unpaired.

**Figure S2. Transcriptomic analysis reveals hormonal and reproductive pathways affected by Apelin dysregulation. (A, B)** Volcano plots of downregulated (red) and upregulated (green) genes identified by GSEA in WT_4YO vs. Apelin_Mut_1YO (A) and WT_1YO vs. WT_4YO (B) comparisons. **(C, D)** GO (Gene Ontology) brain aging-associated (C) and aging-independent (D) identified by GSEA analysis. **(E, F)** Enrichment plot network from the brain aging-associated (E) and aging-independent (F) GO analysis.

**Figure S3. Atoh8 is a downstream effector of Apelin signaling. (A, B)** Gene Ontology (GO) identified by GSEA in *apln* loss-(A) or gain-(B) of-function larvae at 6 dpf. **(C)** *In situ* hybridization of *atoh8* expression in embryos injected with *apln* mRNA and fixed at 24 hpf. **(D, E, F)** Distribution of phenotypes in uninjected control fish (n=200) (D), *apln* +/- incross (n= 468) (E), and control fish injected with *atoh8* mRNA (n=182) (F)**. (G, H)** Distribution of phenotypes and genotypes in *apln* +/- incross (n= 872) (G) and *apln* +/- incross injected with *atoh8* mRNA (n=431) (H)**. (I, J)** Phenotypes followed from 1 dpf (inner ring) to 3 dpf (outer ring) in uninjected control fish (n= 141) (I) and fish injected with *atoh8* CRISPR gRNA + CAS-9 (n= 137) (J). Phenotypes and genotypes of single larva can be visualized as follows: dead (red), severe (orange), mild (yellow), normal (green), wild-type (light grey), heterozygous (medium grey) and homozygous (dark grey). Inner ring: global genotype distribution (%) across the sampled population. Outer ring: genotype frequencies (%) within each phenotype subpopulation. **(K)** *atoh8* gene sequence in control (top) and *atoh8* CRISPR-injected fish (bottom). **(L)** Mutation efficiency in individual larvae after *atoh8* gRNA + CAS-9 injection: wild-type genotype (n= 1) and mutated genotype (n= 23). **(M)** Representative images of phenotypes observed in F0 *atoh8* gRNA larvae at 3 dpf. Scale bars: 100 μm.

## Notes

### Competing Interest Statement

The authors have declared no competing interest.

