## Supplementary material for "Apelin signaling controls neuroendocrine homeostasis via Atoh8-dependent transcriptional regulation": SuppFigures

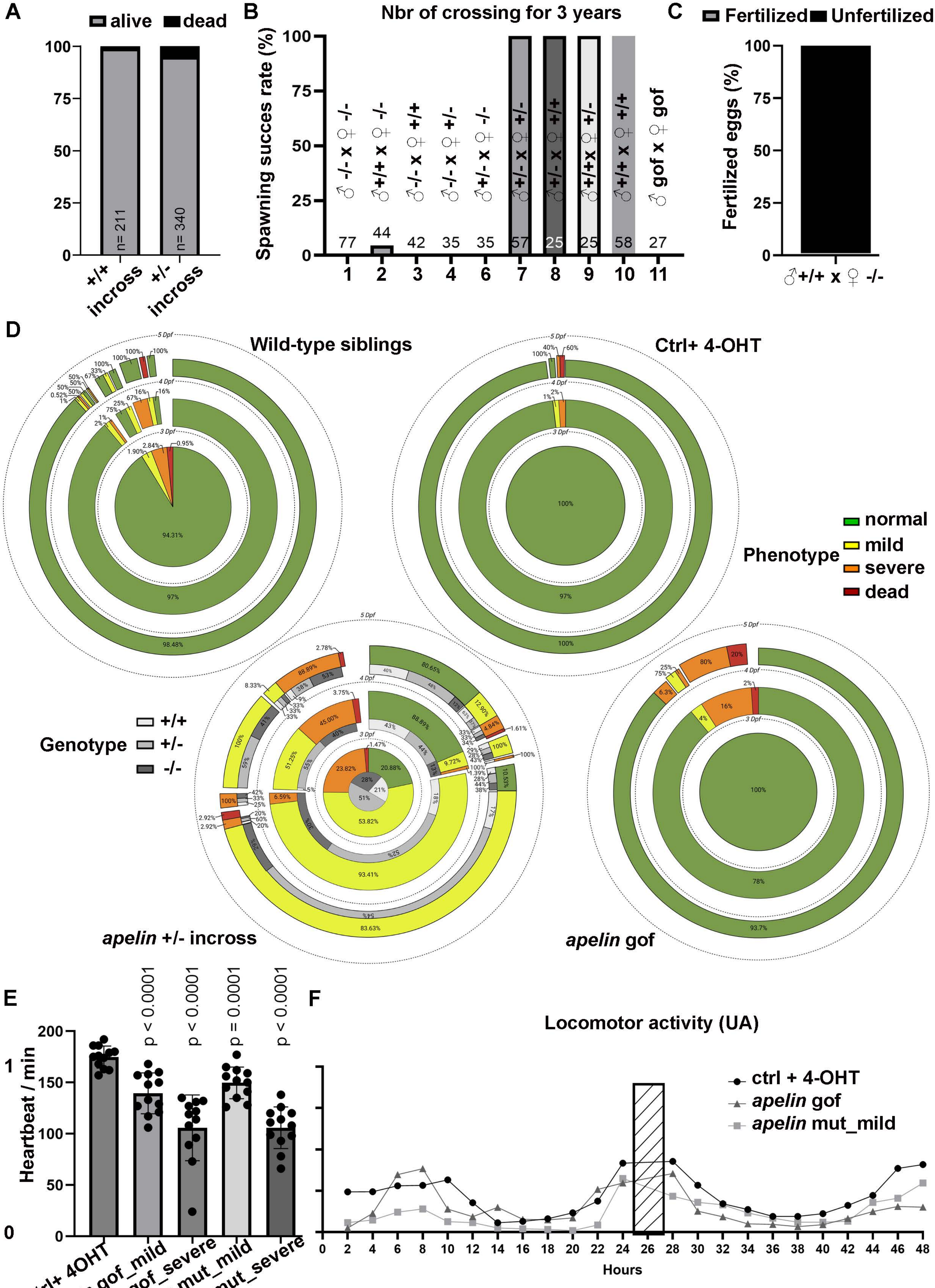
A
alive
dead
100
75
50
25
0
n= 211
n= 340
+/+ incross
+/- incross
B
Nbr of crossing for 3 years
Spawning succes rate (%)
♂-/- x ♀-/-
♂+/+ x ♀-/-
♂-/- x ♀+/+
♂-/- x ♀+/-
♂+/- x ♀-/-
♂+/- x ♀+/-
♂+/- x ♀+/+
♂+/+x ♀+/-
♂+/+ x ♀+/+
♂ gof x ♀ gof
77
44
42
35
35
57
25
25
58
27
1
2
3
4
6
7
8
9
10
11
C
Fertilized
Unfertilized
Fertilized eggs (%)
♂+/+ x ♀ -/-
D
Wild-type siblings
Ctrl+ 4-OHT
5 Dpf
4 Dpf
3 Dpf
94.31%
97%
98.48%
100%
97%
100%
Phenotype
normal
mild
severe
dead
Genotype
+/+
+/-
-/-
apelin +/- incross
53.82%
93.41%
83.63%
80.65%
88.89%
45.00%
51.25%
apelin gof
100%
78%
93.7%
80%
20%
16%
E
Heartbeat / min
200
150
100
50
0
p < 0.0001
p < 0.0001
p = 0.0001
p < 0.0001
ctrl+ 4OHT
apln gof_mild
apln gof_severe
apln mut_mild
apln mut_severe
F
Locomotor activity (UA)
ctrl + 4-OHT
apelin gof
apelin mut_mild
Hours
0
2
4
6
8
10
12
14
16
18
20
22
24
26
28
30
32
34
36
38
40
42
44
46
48

Figure S1

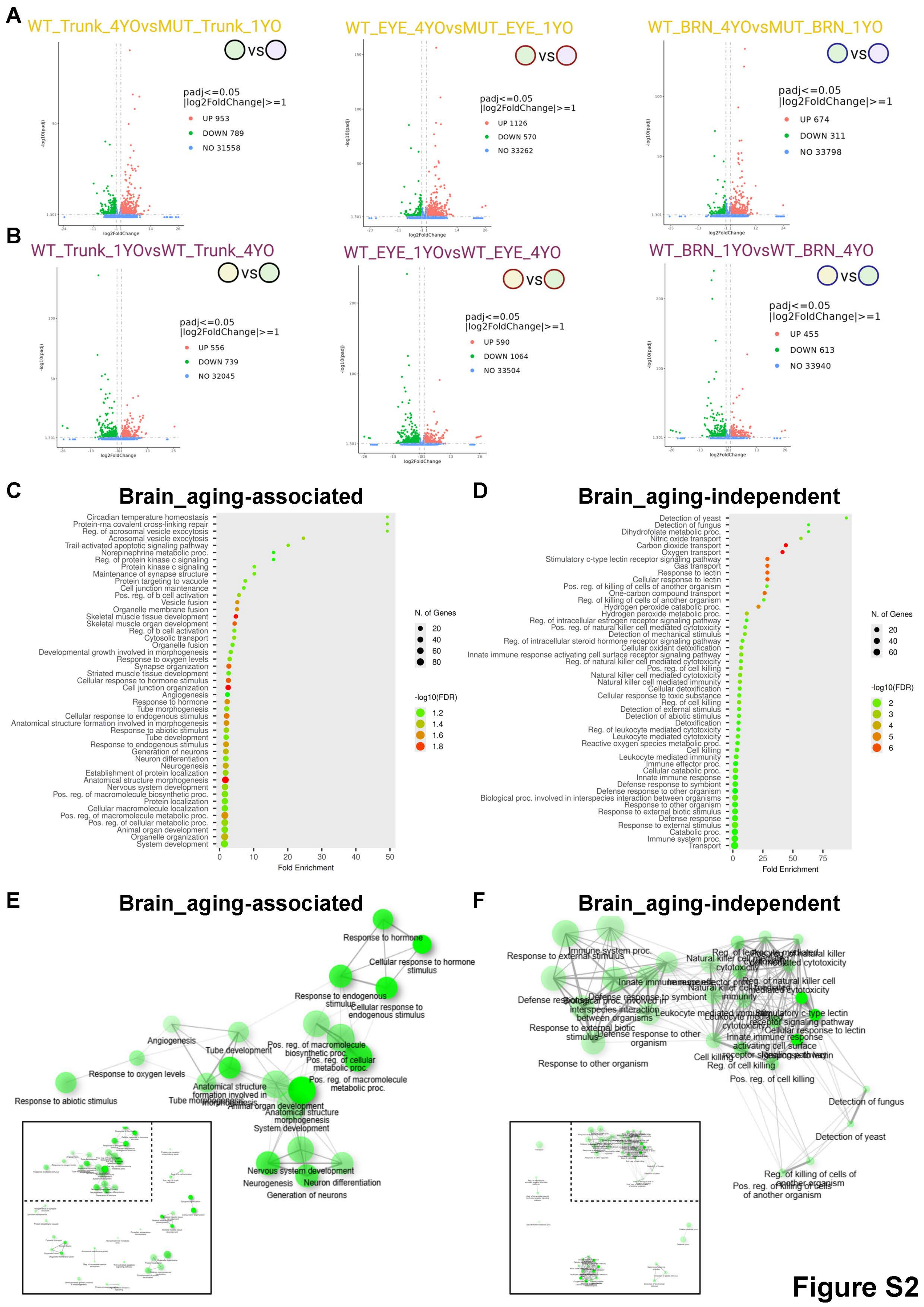
A
WT_Trunk_4YOvsMUT_Trunk_1YO
WT_EYE_4YOvsMUT_EYE_1YO
WT_BRN_4YOvsMUT_BRN_1YO
padj<=0.05
|log2FoldChange|>=1
UP 953
DOWN 789
NO 31558
UP 1126
DOWN 570
NO 33262
UP 674
DOWN 311
NO 33798
B
WT_Trunk_1YOvsWT_Trunk_4YO
WT_EYE_1YOvsWT_EYE_4YO
WT_BRN_1YOvsWT_BRN_4YO
UP 556
DOWN 739
NO 32045
UP 590
DOWN 1064
NO 33504
UP 455
DOWN 613
NO 33940
C
Brain_aging-associated
D
Brain_aging-independent
N. of Genes
-log10(FDR)
Fold Enrichment
E
Brain_aging-associated
F
Brain_aging-independent
Figure S2

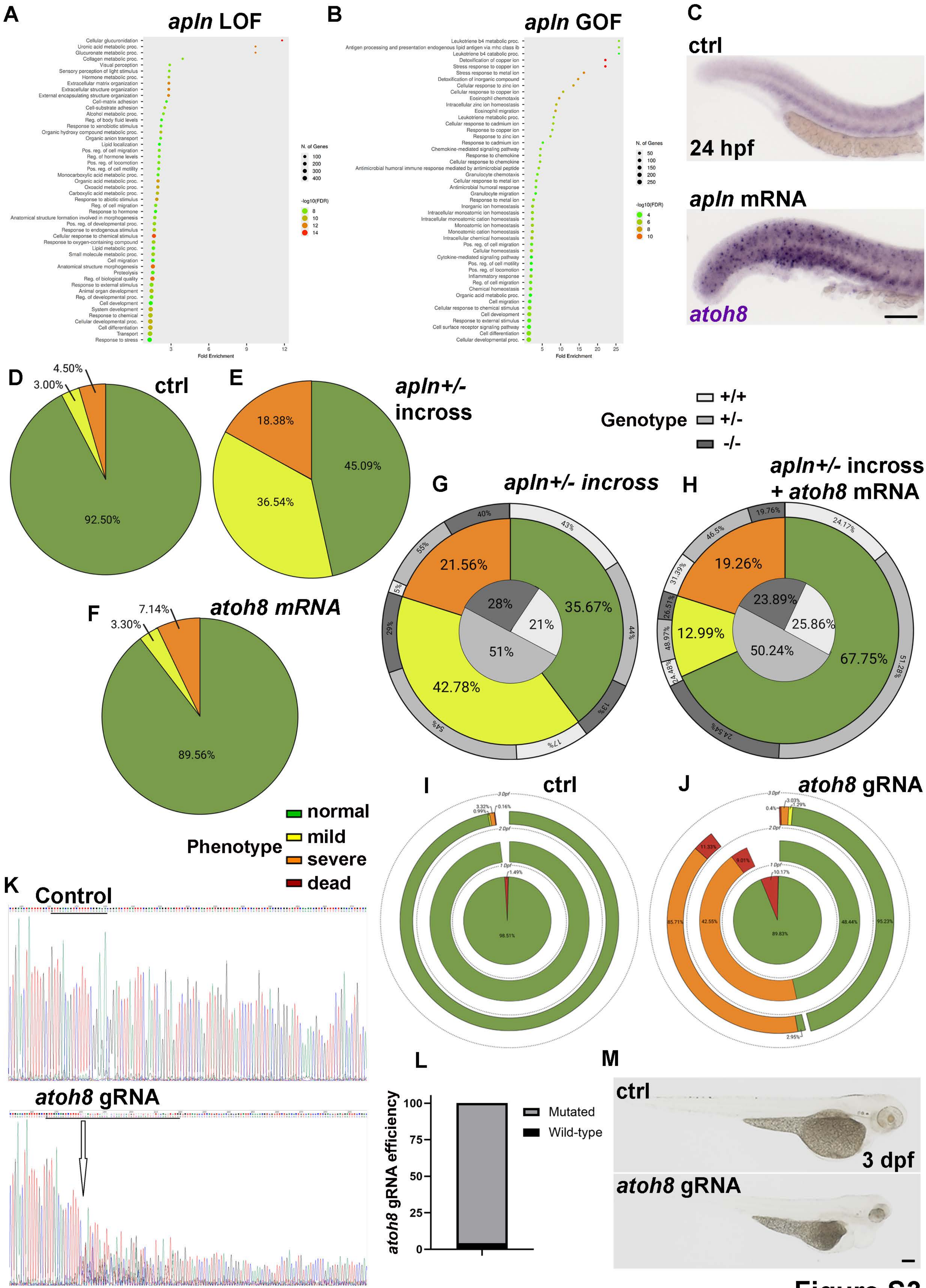

A
apln LOF
B
apln GOF
C
ctrl
24 hpf
apln mRNA
atoh8
D
ctrl
92.50%
3.00%
4.50%
E
apln+/- incross
18.38%
36.54%
45.09%
Genotype
+/+
+/-
-/-
G
apln+/- incross
21.56%
42.78%
35.67%
28%
21%
51%
H
apln+/- incross + atoh8 mRNA
19.26%
12.99%
67.75%
23.89%
25.86%
50.24%
F
atoh8 mRNA
7.14%
3.30%
89.56%
Phenotype
normal
mild
severe
dead
I
ctrl
J
atoh8 gRNA
K
Control
atoh8 gRNA
L
atoh8 gRNA efficiency
Mutated
Wild-type
M
ctrl
3 dpf
atoh8 gRNA

Figure S3
